# Experimental considerations for study of *C. elegans* lysosomal proteins

**DOI:** 10.1101/2022.06.30.498309

**Authors:** John C. Clancy, An A. Vo, Krista M. Myles, Max T. Levenson, James Matthew Ragle, Jordan D. Ward

## Abstract

Lysosomes are an important organelle required for the degradation of a range of cellular components. Lysosome function is critical for development and homeostasis as dysfunction can lead to inherited genetic disorders, cancer, and neurodegenerative and metabolic disease. The acidic and protease-rich environment of lysosomes poses experimental challenges. Many fluorescent proteins are quenched or degraded, while specific red fluorescent proteins can be cleaved from translational fusion partners and accumulate. While studying MLT-11, a *C. elegans* molting factor that localizes to lysosomes and the cuticle, we sought to optimize several experimental parameters. We found that, in contrast to mNeonGreen fusions, mScarlet fusions to MLT-11 missed cuticular and rectal epithelial localization. Rapid sample lysis and denaturation was critical for preventing MLT-11 fragmentation while preparing lysates for western blots. Using a model lysosomal substrate (NUC-1) we found that rigid polyproline linkers and truncated mCherry constructs do not prevent cleavage of mCherry from NUC-1. We provide evidence that extended localization in lysosomal environments prevents the detection of FLAG epitopes in western blots. Finally, we optimize an acid-tolerant green fluorescent protein (Gamillus) for use in *C. elegans*. These experiments provide important experimental considerations and new reagents for the study of *C. elegans* lysosomal proteins.

## INTRODUCTION

Lysosomes are membrane-enclosed cytoplasmic organelles required for the degradation of diverse biological macromolecules (Ballabio and Bonifacino 2020). Consistent with this function, they are among the most acidic compartment in the cell with a pH ranging from 4.5-5.5, and are packed with proteases, nucleases, acid lipases, and carbohydrate processing enzymes (Bonam *et al*. 2019). Lysosome dysfunction can lead to inherited lysosomal storage disorders, as well as neurodegenerative and metabolic disease, and cancer (Ballabio and Bonifacino 2020). Lysosome activity declines with age and is required for lifespan extension (Hansen *et al*. 2008; Sun *et al*. 2020).

Using fluorescent protein (FP) fusions to study lysosomal lumen proteins present challenges. Many green and red FPs derived from avGFP and eqFP578, respectively, are sensitive to degradative lysosomal proteases (Shinoda *et al*. 2018b). The sensitivity of many other FPs to lysosomal proteases remains to be determined (Shinoda *et al*. 2018b). Due to their low pKa (3.1-5.3) and resistance to lysosomal proteases, red FPs derived from DsRed or eqFP611 (i.e. mCherry, mScarlet, mRuby) are typically the FP of choice for imaging lysosomal lumen proteins (Shinoda *et al*. 2018b). An additional consideration in interpreting lysosomal localization is that lysosomal proteases can cleave flexible linkers or the N-terminus of FPs, separating the FP from the protein of interest (Ko *et al*. 2003; Kollmann *et al*. 2005; Huang *et al*. 2014; Miao *et al*. 2020). While this cleavage can be used to monitor lysosomal activity (Miao *et al*. 2020), it can hamper interpretation of lysosomal localization of fusion proteins. Another issue is that many FPs lose fluorescence in the acidic lysosome through fluorophore quenching due to their neutral pKa (Shinoda *et al*. 2018b). Acid-tolerant green FPs have been recently developed but have not yet been widely adopted (Roberts *et al*. 2016; Shinoda *et al*. 2018b).

During the course of studying MLT-11, a putative *C. elegans* protease inhibitor, we used CRISPR/Cas9 to introduce an *mScarlet∷3xMyc* tag into the endogenous *mlt-11* locus to produce a C-terminal translational fusion that should label all isoforms (Ragle et al. 2022). This strain displayed robust MLT-11∷mScarlet∷3xMyc localization in punctae and tubules reminiscent of lysosomes. However, we were unable to verify the fusion was full-length by anti-Myc western blotting. We also generated an equivalent MLT-11∷mNeonGreen∷3xFLAG fusion without a linker (Ragle et al. 2022), which displayed similar punctate/tubular localization, but also transient cuticular localization. This discrepancy between these strains motivated us to explore whether we could minimize cleavage of the fluorescent protein fusion and explore acid-tolerant green FPs for lysosomal translational fusions.

## MATERIALS AND METHODS

### Strains and culture

*C. elegans* were cultured as originally described (Brenner 1974), except worms were grown on MYOB media instead of NGM. MYOB agar was made as previously described (Church *et al*. 1995). We obtained wild type N2 animals from the *Caenorhabditis* Genetics Center (CGC).

### Strains used in this study

Strains used in this study:
| Strain | Construction/Source | Genotype |
| --- | --- | --- |
| N2 | CGC | Wild type |
| JDW206 | CRISPR/Cas9 | *mlt-11(wrd41[mlt-11::30x linker-mScarlet-I<sup>Δ</sup>3xMyc])* |
| JDW254 | Crossing | *qxIs630[scav-3::GFP] ; mlt-11(wrd41[mlt-11::30 amino acid linker-mScarlet-I-3xMyc]) V* |
| JDW372 | CRISPR/Cas9 | *his-72(wrd75[his-72::Gamillus] III* |
| JDW373 | CRISPR/Cas9 | *lmn-1(wrd76[lmn-1::Gamillus]) I* |
| JDW376 | RMCE | *jsTi1493 {mosL::loxP::mex-5p::FLP::sl2::mNeonGreen::rpl-28p::FRT [wrdSi67(SEC::loxP::mlt-11p (-2.8 kb)::nuc-1::mCherry-tbb-2 3'UTR)] FRT3::mosR} IV* |
| JDW379 | SEC excision of JDW376 | *jsTi1493 {mosL loxP [wrdSi70(mlt-11p (-2.8 kb)::nuc-1::mCherry-tbb-2 3'UTR)] FRT3::mosR} IV* |
| JDW381 | RMCE | *jsTi1493 {mosL::loxP::mex-5p::FLP::sl2::mNeonGreen::rpl-28p::FRT [wrdSi71(SEC::loxP::mlt-11p (-2.8 kb)::nuc-* |
|  |  | *1::P5::crmCherry::3xFLAG-tbb-2 3'UTR)]*<br>*FRT3::mosR} IV* |
| JDW382 | SEC excision of JDW381 | *jsTi1493 {mosL loxP [wrdSi71(mlt-11p (-2.8 kb)::nuc-1::P5::crmCherry::3xFLAG-tbb-2 3'UTR)]*<br>*FRT3::mosR} IV* |
| JDW391 | Ragle et al., 2022 | *mlt-11(wrd86[C-terminal mNeonGreen::3xFLAG])*<br>*V* |
| JDW499 | Rapid RMCE | *jsSi1579 jsSi1706 jsSi1726 wrdSi90[loxP myo-2p::NLS::mNeonGreen, rps-0p HygR, loxP hsp-16.41p::nuc-1::P5 linker::crmCherry::3xFLAG::tbb-2 3'UTR FRT3]* |
| JDW382 | RMCE | *jsTi1493 {mosL loxP [wrdSi71(mlt-11p (-2.8 kb)::nuc-1::P5::crmCherry::3xFLAG-tbb-2 3'UTR)]*<br>*FRT3::mosR} IV* |
| JDW539 | CRISPR/Cas9 | *glh-1(wrd151[glh-1::Gamillus]) I* |
| JDW556 | RMCE | *jsSi1579 jsSi1706 jsSi1726 wrdSi106[loxP myo-2p::NLS::mNeonGreen, rps-0p HygR, loxP hsp-16.41p::nuc-1::linker::Gamillus::linker::mCherry::tbb-2 3'UTR FRT3]* |
| JDW557 | RMCE | *jsSi1579 jsSi1706 jsSi1726 wrdSi107[loxP myo-2p::NLS::mNeonGreen, rps-0p HygR, loxP hsp-16.41p::nuc-1::linker::Gamillus::linker::mScarlet-I::tbb-2 3'UTR FRT3]* |
| NM5179 | CGC | *jsTi1493 [mosL loxP mex-5p FLP sl2 mNeonGreen rpl-28p FRT GFP-HIS-58 FRT3 mosR] IV* |
| NM5548 | CGC | *jsSi1579 jsSi1706 jsSi1726[loxP myo-2p FRT nlsCyOFP myo-2 3' mex-5p FLP D5 glh-2 3' FRT3] II* |
| XW8056 | CGC | *qxIs630[scav-3::GFP]* |
| XW13734 | Dr. Xiaochen Wang | *PhspNUC-1::sfGFP::mCherry* |

### Transgenesis and genome editing

All plasmids used are listed in Table S1. Annotated plasmid sequence files are provided in File S1. Specific cloning details and primers used are available upon request. An *hsp-16.41∷linker∷Gamillus∷linker∷mCherry∷tbb-2 3’UTR* cassette was synthesized and cloned (Twist Bioscience) to create pJW2138 (Table S1). *nuc-1* coding sequence was Gibson cloned into this vector to create pJW2139 (Table. S1; *hsp-16.41∷nuc-1∷linker∷Gamillus∷linker∷mCherry∷tbb-2 3’UTR*). The mCherry cassette in pJW2139 was replaced with mScarlet-I through Gibson cloning to create pJW2145. The *hsp-16.41∷nuc-1∷linker∷Gamillus∷linker∷red FP∷tbb-2 3’UTR* cassettes from pJW2139 and pJW2145 were PCR amplified and SapTrap cloned (Schwartz and Jorgensen 2016) into pNM4216 to generate pJW2460 and pJW2461, respectively. pNM4216 is an insertion vector for rapid recombination mediated cassette exchange (rRMCE). rRMCE is a derivative of RMCE which generates usable knock-ins more quickly by removing the need to excise a selectable marker (see https://sites.wustl.edu/nonetlab/rapid-rmce-beta-testing/ last updated 6-15-2022 for more details and for protocols). pJW2460 and pJW2461 were integrated into NM5548 using rRMCE to generate strains JDW556 and JDW557, respectively.

We used Q5 site-directed mutagenesis (NEB) on pJW2139 to truncate mCherry, remove Gamillus and replace the linker with a rigid penta-proline linker to generate pJW2201. A 3xFLAG tag was added by Gibson cloning to create pJW2204 (Table S1; *hsp-16.41p∷nuc-1∷P5 linker∷crmCherry∷tbb-2 3’UTR*). The *nuc-1∷P5 linker∷crmCherry∷3xFLAG∷tbb-2 3’UTR* cassette was PCR amplified and ATG and GTA connectors for SapTrap cloning were added (Schwartz and Jorgensen 2016). This PCR product was Gibson cloned to create pJW2325. *nuc-1* and *linker∷mCherry∷tbb-2 3’UTR* fragments were PCR amplified from pJW2139 and ATG and GTA SapTrap connectors were added (Schwartz and Jorgensen 2016). These products were Gibson cloned to create pJW2322. A 2.8 kb *mlt-11* promoter fragment PCR amplified with SapTrap TGG and ATG connectors and Gibson cloned to generate pJW2286. Integration vectors (pJW2328 and pJW2331) for standard RMCE were created by SapTrap with a pLF3FShC backbone (Schwartz and Jorgensen 2016; Nonet 2020). JDW379 and JDW382 were created by RMCE using strain NM5179 and pJW2328 and pJW2331, respectively (Nonet 2020).

JDW206 was created using CRISPR/Cas9-mediated genome editing with a pJW1897 repair template and a pJW1896 sgRNA plasmid. Knock-ins were generated, and the self-excising cassette was excised as previously described (Dickinson *et al*. 2015). pJW1896 was created by SapTrap using a pJW1839 backbone (Schwartz and Jorgensen 2016; Ashley et al. 2021). pJW1897 was created by SapTrap with 600 bp 5’ and 3’ homology arms and a pJW1821 (30 amino acid linker∷mScarlet (GLO)^SEC Lox511I^3xMyc) cassette (Schwartz and Jorgensen 2016; Ashley *et al*. 2021). Gamillus knock-ins into *glh-1, his-72 and lmn-1* and *GFP* knock-ins into *glh-1* were generated by injection of Cas9 ribonucleoprotein complexes [700 ng/ul IDT Cas9, 115 ng/ul crRNA and 250 ng/ul IDT tracrRNA] and a dsDNA repair template (25-50 ng/ul) created by PCR amplification of a plasmid template (Paix et al., 2014; Paix et al., 2015). The PCR products were melted to boost editing efficiency, as previously described (Ghanta and Mello, 2020). crRNAs used are provided in Table S2. Oligonucleotides used for repair template generation from template pJW2139 or pJW2088 (Ashley et al., 2021) and for genotyping are provided in Table S3.

### Microscopy

Animals were picked into a 5 μl drop of M9 + 0.05% with levamisole solution on a 2% agarose pad on a microscope slide, then a coverslip was placed on the pad. Images were acquired using a Plan-Apochromat 40x/1.3 Oil DIC lens or a Plan-Apochromat 63x/1.4 Oil DIC lens on an AxioImager M2 microscope (Carl Zeiss Microscopy, LLC) equipped with a Colibri 7 LED light source and an Axiocam 506 mono camera. Acquired images were processed through Fiji software (version: 2.0.0-rc-69/1.52p) (Schindelin *et al*. 2012). For direct comparisons within a figure, we set the exposure conditions to avoid pixel saturation of the brightest sample and kept equivalent exposure for imaging of the other samples. For co-localization analysis, animals of the indicated genotype were synchronized by alkaline bleaching (dx.doi.org/10.17504/protocols.io.j8nlkkyxdl5r/v1), released on MYOB plates, and incubated at 20°C for 48 hours. Plates were heat-shocked at 34°C for 30 minutes and then incubated at 20°C for an additional 24 hours. Animals were picked and imaged with a 63X objective, as described above except no agarose pad was used for image acquisition. Consistent exposure times for green and red FP imaging were used for each strain. Background was removed using a rolling ball method in Fiji (radius=50). Subsequent analyses were performed using Imaris software (Oxford Instruments). A mask was created using surface detail=10 microns, voxel intensity=10. The Coloc tool was then used with a threshold set to 0.25 and PSF width set to 0.25. A Mander’s test was performed on the colocalization data (Manders *et al*. 1992, 1993).

### Western blotting

For the western blot in Fig. 2, JDW391 animals were synchronized by alkaline bleaching (dx.doi.org/10.17504/protocols.io.j8nlkkyxdl5r/v1) and released on MYOB plates. Animals were harvested at 42 hours post-release by picking thirty animals into 30 μl of M9+0.05% gelatin. Samples were processed as described in Fig 2A. For all other western blots, forty animals were picked into 40 μl of M9+0.05% gelatin and Laemmli sample buffer was added to 1X and then immediately incubated for five minutes at 95°C. Lysates were then stored at −80°C until they were resolved by SDS-PAGE. For the western blots in Figure 3, animals were synchronized by bleaching and harvested at the indicated times. Lysates were resolved using precast 4-20% MiniProtean TGX Stain Free Gels (Bio-Rad) with a Spectra™ Multicolor Broad Range Protein Ladder (Thermo; # 26623) protein standard. Proteins were transferred to a polyvinylidene difluoride membrane by semi-dry transfer with a TransBlot Turbo (Bio-Rad). Blots and washes were performed as previously described. Anti-FLAG blots used horseradish peroxidase (HRP) conjugated anti-FLAG M2 (Sigma-Aldrich, A8592-5×1MG, Lot #SLCB9703) at a 1:2000 dilution.

Mouse anti-alpha-Tubulin 12G10 (Developmental Studies Hybridoma Bank; “-c” concentrated supernatant) was used at 1:4000. Rabbit anti-mCherry (AbCam ab167453) was used at 1:1000. The secondary antibodies were Digital anti-mouse (Kindle Biosciences LLC, R1005) diluted 1:20,000 or Digital anti-Rabbit (Kindle Biosciences LLC, R1006) diluted 1:1000. Blots were incubated for 5 minutes with 1 ml of Supersignal West Femto Maximum Sensitivity Substrate (Thermo Fisher Scientific, 34095) and the final blot were imaged using the ‘chemi high-resolution’ setting on a Bio-Rad ChemiDoc MP System.

## Data availability

Strains and plasmids are available upon request. To facilitate generation of repair templates and subcloning, all plasmid sequences are provided in File S1, knock-in sequences are provided in File S2. Sequences of oligonucleotides used for cloning are available upon request.

## RESULTS

### MLT-11∷mScarlet localizes to lysosomes and the vulva but not the aECM or rectal lining

We recently demonstrated that *C. elegans* MLT-11 is secreted protein that localizes in the cuticle and in a punctate pattern reminiscent of lysosomes (Ragle et al. 2022). The cuticular expression in L4 is transient which might suggest that it oscillates, though further work is required to test this assertion (Ragle et al,.2022). Our initial attempts at generating a MLT-11 translational reporter involved inserting an mScarlet∷3xMyc cassette with a flexible 30 amino acid linker to C-terminally tag all known *mlt-11* isoforms (Fig. 1A). This knock-in displayed vulval localization (Fig. 1B) similar to the MLT-11∷mNeonGreen∷3xFLAG (MLT-11∷mNG) fusion (Ragle et al. 2022). While we observed MLT-11∷mScarlet in rectal epithelial cells (Fig. 1C), we did not observe it lining the rectum as we did for the MLT-11∷mNG fusion (Ragle et al. 2022). There was robust MLT-11∷mScarlet expression in the hypodermis with a range of expression patterns ranging from punctate to tubular (Fig 1D). This pattern resembled NUC-1∷mCherry expression (Miao *et al*. 2020), suggesting MLT-11:mScarlet might localize to lysosomes. Accordingly, MLT-11∷mScarlet co-localized with the lysosomal marker, SCAV-3∷GFP (Fig 1D). We observed a similar expression pattern for MLT-11∷mNG (Fig. 1E), but also cuticular expression that was not observed for MLT-11∷Scarlet (Fig. 1D, E). These data highlight that mNG and mScarlet fusions to equivalent positions in a protein can produce different localization patterns.

**Fig. 1.**
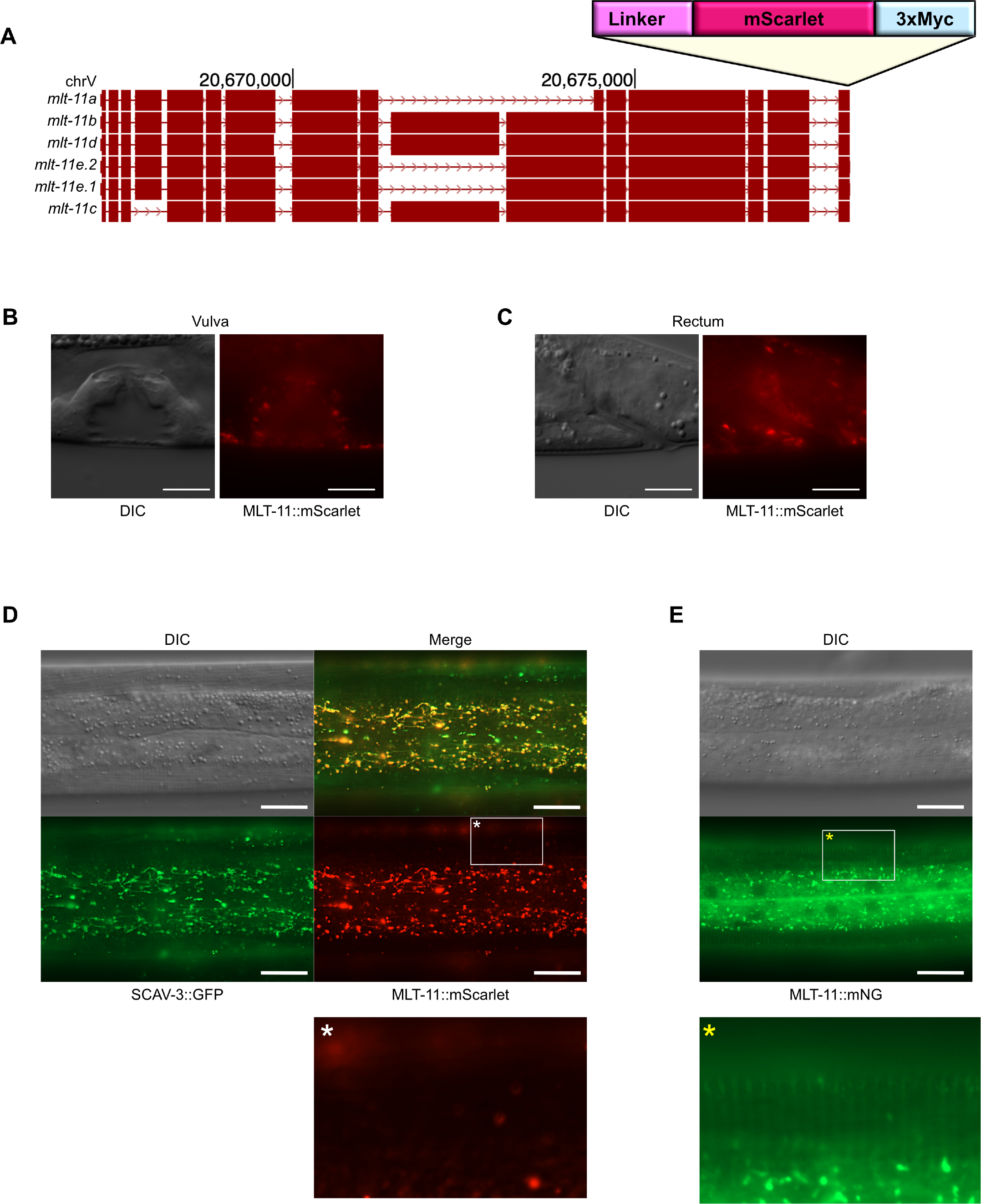
MLT-11∷mScarlet is detected in lysosomes but not the cuticle. (A) Schematic from the UCSC genome browser of the *mlt-11* locus and isoforms. Position in bp on chromosome V (chrV) is indicated. The insertion site of the *linker∷mScarlet∷3xMyc* tag is indicated. DIC and MLT-11∷mScarlet images of L4.6 animals depicting localization in the vulva (B) and rectum (C). Images are representative of 50 animals imaged over 3 independent experiments. Scale bars=10 μm. (D) Representative images of *mlt-11∷mScarlet∷3xMyc; scav-3∷GFP* animals. SCAV-3 is a lysosomal membrane protein used as a marker for lysosomal localization (Li et al., 2016). (E) Representative images of *mlt-11∷mNeonGreen∷3xFLAG* animals. In (D and E) scale bars=20 μm. Images are representative of >100 animals examined over four independent experimentsAsterisks indicate a region of the cuticle that is provided as a zoomed in image below.

### Sample processing affects MLT-11∷mNG stability in lysate generation

As fluorescent tags can be cleaved off fusion proteins in lysosomes (Miao *et al*. 2020), western blotting to confirm that a fusion protein is full-length is essential to have high confidence in lysosomal localization. For our *mlt-11∷mScarlet∷3xMyc* strain we were never able to detect bands of the predicted size by western blotting with anti-Myc or anti-mScarlet antibodies (unpublished data). When attempting western blots on *mlt-11∷mNeonGreen∷3xFLAG* lysates, we saw variable laddering (for example see Fig 2B, lane C). As lysosomal proteases can degrade proteins, we sought to optimize our sample preparation conditions to minimize degradation. We harvested samples at peak MLT-11 protein expression (42 hours post-release, stage L4.3 (Mok *et al*. 2015; Ragle *et al*. 2022), and tested a range of variables: i) denaturation at 70°C for 10 minutes vs. 95°C for 5 minutes ii) denaturing samples immediately after collection vs. rapid freezing and denaturation of all samples together later; iii) rapid freezing using dry ice vs. liquid nitrogen; and iv) whether it was better to denature before storage at −80°C vs denature immediately before resolving samples by SDS-PAGE (Fig. 2A). The best approach was to harvest animals, add Laemmli sample buffer and immediately denature before storage at −80°C (Fig. 2B lane D and H). Denaturation at 95°C for 5 minutes produced less laddering than heating to 70°C for 10 minutes (Fig 2B compare D to H). The other approaches with various combinations of rapid freezing and denaturation all produced more degradation products above 50 kDa (Fig. 2B). In all conditions there is a strong band at 50 kDa (Fig. 2B), consistent with a C-terminal MLT-11 fragment we previously observed (Ragle et al., 2022). These experiments demonstrate that sample preparation has a significant effect on MLT-11 stability during preparation of lysates for immunoblotting.

**Fig. 2.**
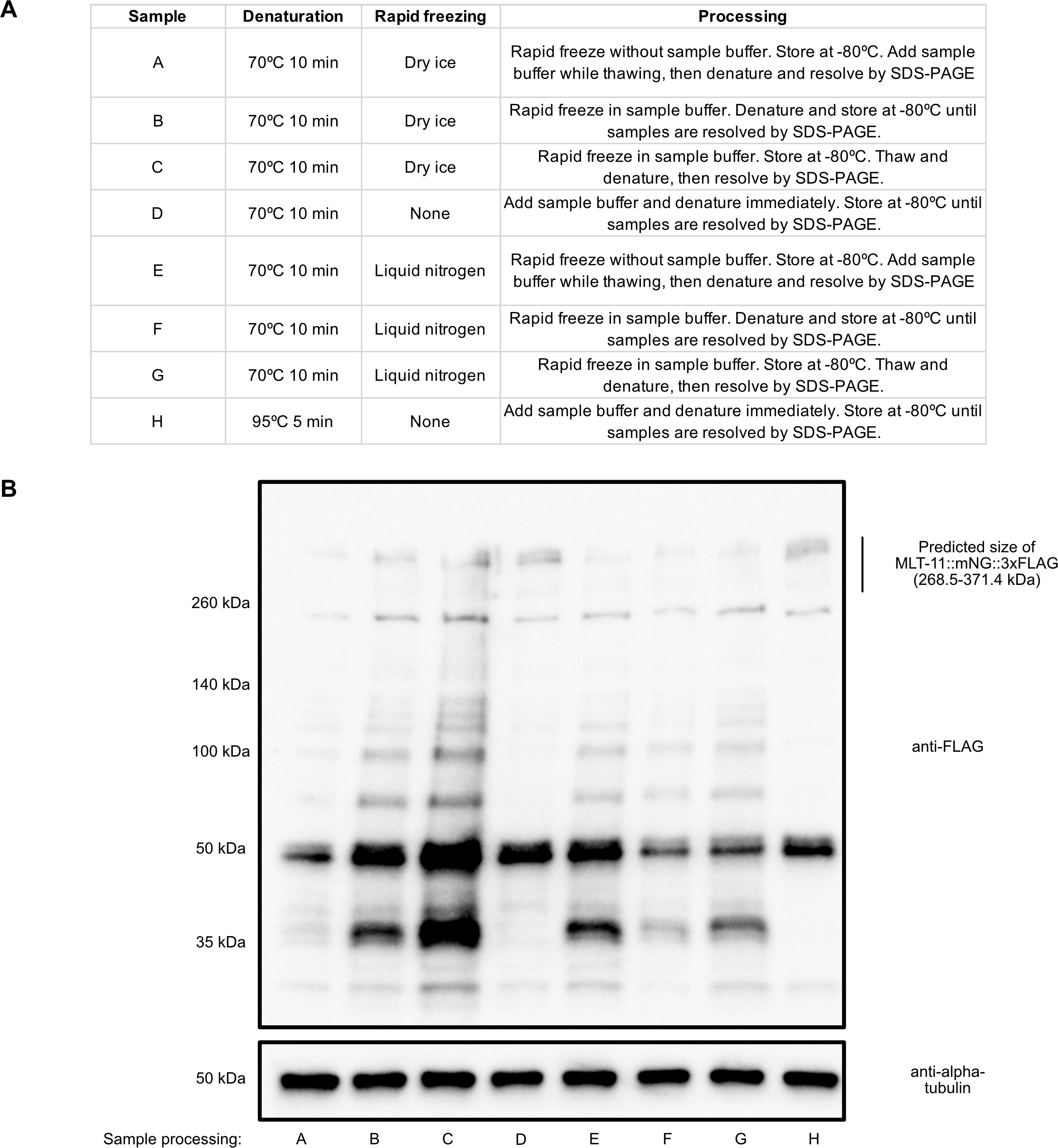
Immediate denaturation of *mlt-11∷mNeonGreen∷3xFLAG* samples minimizes degradation. (A) Table describing different sample processing regimens. (B) Anti-FLAG and anti-alpha-tubulin immunoblots on *mlt-11∷mNeonGreen∷3xFLAG* lysates processed using the conditions described in A. Blot is a representative of two independent experiments. Predicted size of full-length MLT-11∷mNeonGreen∷3xFLAG is provided in kilodaltons (kDa).

### Tag cleavage is not reduced by proline linkers or truncated mCherry

As red FPs are stable in lysosomes and linkers can be cleaved by lysosomal proteases (Shinoda *et al*. 2018b), red FP accumulation might not reflect true localization of a fusion protein. We therefore tested whether we could design FP fusions that underwent minimal cleavage. We used a well-characterized *nuc-1∷mCherry* translational fusion as our test case, expressing it in hypodermal and seam cells with a strong *mlt-11* promoter (Ragle et al. 2022). NUC-1∷mCherry is cleaved by lysosomal proteases and this cleavage is more frequent when lysosomes acidify during molting (Miao *et al*. 2020). In mammalian cells, rigid linkers comprised of five prolines (P5) helps minimize lysosomal cleavage, as does removing the eleven N-terminal amino acids of mCherry to make a cleavage resistant version (crmCherry)(Huang *et al*. 2014). To test whether these modifications reduce NUC-1∷mCherry cleavage in *C. elegans* lysosomes, we generated *mlt-11p∷nuc-1∷P5∷crmCherry∷3xFLAG* single-copy transgenes. We also generated a *mlt-11p∷nuc-1∷mCherry* strain with the equivalent linker (GGGSRGGTR) used in the *nuc-1∷mCherry* constructs of Miao *et al*. (2020). We harvested synchronized mid-L4 larvae, late-L4 larvae, and adults for imaging. For both strains, we observed robust lysosomal mCherry expression at all timepoints (Fig. 3B). We also collected animals for western blot analysis. NUC-1∷mCherry and NUC-1∷P5∷crmCherry∷3xFLAG displayed similar punctate and tubular localization at each timepoint (Fig. 3B). We observed similar cleavage levels of NUC-1∷mCherry and NUC-1∷P5∷crmCherry∷3xFLAG, suggesting that the P5 linker and N-terminal truncation were not effective at preventing cleavage of the mCherry tag (Fig. 3A). The NUC-1∷mCherry control displayed increased cleavage in late-L4 larvae, similar to previous reports (Miao *et al*. 2020). NUC-1∷P5∷crmCherry∷3xFLAG displayed a unique cleavage product compared to NUC-1∷mCherry, suggesting that the P5 linker, the N-terminal truncation, and/or the FLAG tag were causing cleavage within NUC-1 (Fig. 3A).

**Fig. 3.**
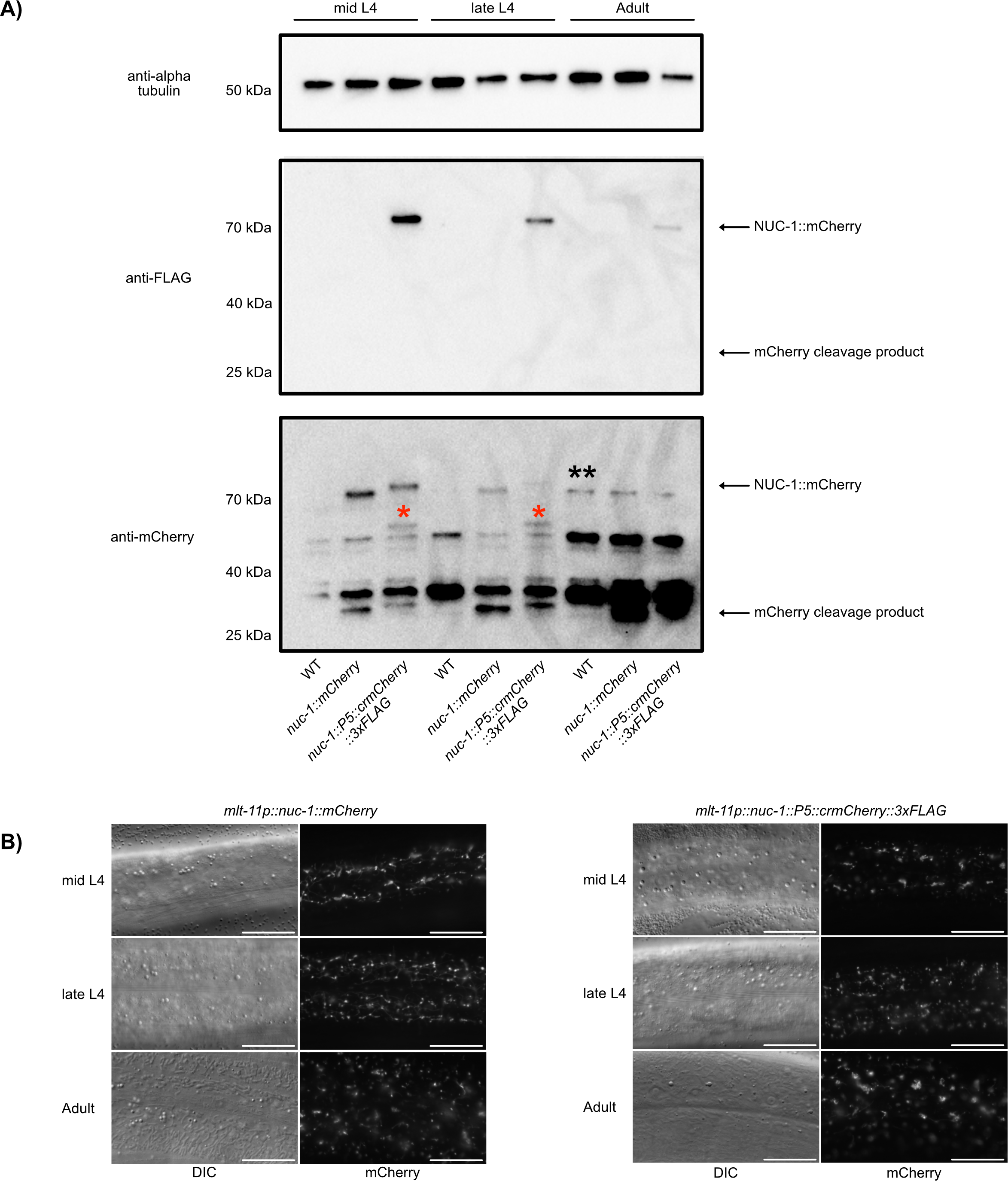
Rigid proline linkers and truncated mCherry does not prevent cleavage from a NUC-1 fusion. Animals of the indicated genotype were synchronized and harvested in mid L4 (48 hours post-release), late L4 (56 hours post-release), and adulthood (72 hours post-release) for immunoblotting (A) and imaging (B). We performed anti-alpha tubulin, anti-FLAG, and anti-mCherry immunoblots on lysates from the indicated genotypes (A). The blots and images are representative of three experimental replicates. Marker size (in kDa) is provided. Full-length NUC-1∷mCherry fusions and cleavage products are indicated by arrows. An mCherry∷3xFLAG specific NUC-1 cleavage product is indicated by red asterisks over the band. A non-specific band only detected in adult lysates is indicated by two black asterisks. (B) Animals of the indicated genotype were imaged at mid L4, late L4, and adulthood. DIC and mCherry images are provided for each strain and time point. Scale bars=20 μm. Images are representative of 50 animals examined per genotype in two independent experiments.

### The NUC-1 FLAG epitope is not recognized in immunoblotting after extended time in the lysosome

In our FLAG immunoblots, the full-length NUC-1∷P5∷crmCherry∷3xFLAG product declined in intensity in late-L4 and adult animals and we did not observe a band at the expected cleavage product position (Fig. 3A). In contrast, in the anti-mCherry immunoblots the cleavage product increased in intensity in late-L4 and adult animals. *mlt-11* mRNA levels oscillate and the promoter shuts off in mid-L4, so we are monitoring NUC-1∷mCherry and NUC-1∷P5∷crmCherry∷3xFLAG produced by the last pulse of gene expression driven by the *mlt-11* promoter (Frand *et al*. 2005; Hendriks *et al*. 2014; Meeuse *et al*. 2020). These data suggest that the FLAG epitope is not recognized in the cleaved mCherry fragment, an important consideration in interpreting anti-FLAG immunoblots.

### The green FP Gamillus is not quenched in C. elegans lysosomes

Another limitation of FP usage in lysosomes is that many green FPs are quenched and degraded (Shinoda *et al*. 2018b). The quenching could produce a false negative for lysosomal expression of a fusion protein. As co-localization studies frequently rely on red and green FPs, we sought alternate green FPs for lysosomal imaging. Two candidates from the literature were pH-tdGFP and Gamillus. pH-tdGFP is an engineered tandem dimer which is acid-tolerant and stable *in vitro* over a pH range from 3.75-8.50 (Roberts *et al*. 2016). However, we did not pursue this green FP as the tandem dimer would make it a large insertion for knock-ins which could decrease editing efficiency. Gamillus is an acid tolerant monomeric green FP developed through directed evolution of a novel green FP from the flower hat jellyfish, *Olindias formosa* (Shinoda *et al*. 2018a). It has a pKa of 3.4 and is reported to have a useful combination of brightness, photostability, and maturation speed (Shinoda *et al*. 2018a). Gamillus is photo-switchable; at its peak excitation wavelength of 504 nm it is switched to an off state which could be reversed by irradiation with 352-388 nm light (Shinoda *et al*. 2018a). Excitation in the 440-480 nm range produced negligible photochromism, potentially due to a higher on-switching rate (Shinoda *et al*. 2018a).

To test whether Gamillus fluoresces in *C. elegans* lysosomes, we created a heat shock-inducible *nuc-1∷mCherry∷Gamillus* transgene. Gamillus and mCherry co-localized in lysosomes 24 hours post-heat shock including in tubular, acidified lysosomes (Fig. 4A,B). This result is in contrast to NUC-1∷sfGFP∷mCherry, where the sfGFP is quenched over time by the acidic lysosomal environment and there is poor co-localization 24 hours post-heat shock (Fig. 4A,B) (Miao *et al*. 2020). As the pKa of mScarlet is higher than that of mCherry (pKa 5.3 vs.3.1), we used this approach to test whether mScarlet is quenched by the lysosomal environment. We constructed a heat shock-inducible *nuc-1∷mScarlet∷Gamillus* and demonstrated that mScarlet and Gamillus also co-localized, suggesting that mScarlet is not quenched or degraded in the lysosome (Fig. 4A,B). Gamillus had a significantly higher co-localization with mCherry or mScarlet by Manders co-efficient in comparison to sfGFP (Fig. 4B). There was no significant difference in Gamillus co-localization with mCherry and mScarlet (Fig. 4B). These data indicate that Gamillus is a suitable green FP tag for lysosome lumenal proteins and that mScarlet is not quenched in lysosomes.

**Fig 4.**
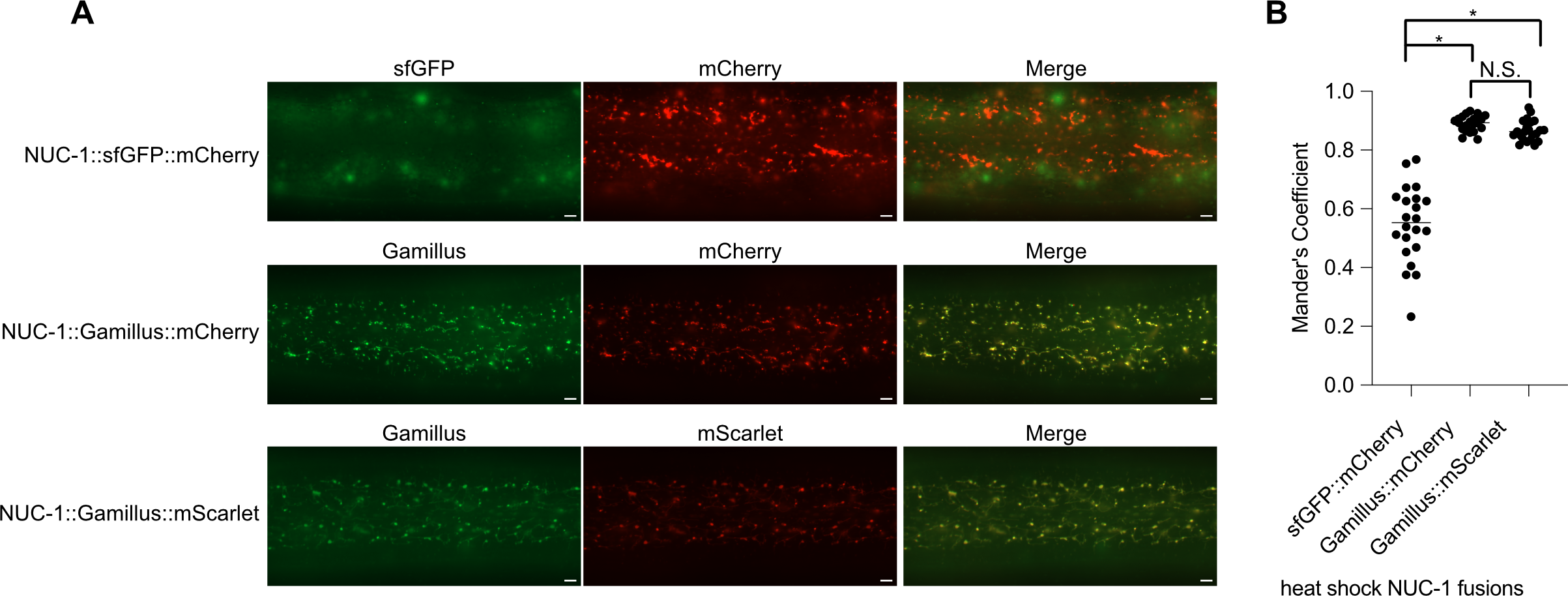
Gamillus is not quenched in lysosomes. *Phsp∷nuc-1∷sfGFP∷mCherry, Phsp∷nuc-1∷Gamillus∷mCherry*, and *Phsp∷nuc-1∷Gamillus∷mScarlet-I* animals were heat-shocked at 34°C for 30 minutes 48 hours post-release and imaged 24 hours post-heat-shock. A representative image for each genotype is provided in (A). Scale bars=5 μm. (B) Mander’s co-localization co-efficient for heat-shock-induced NUC-1∷sfGFP∷mCherry, NUC-1∷Gamillus∷mCherry, and NUC-1∷Gamillus∷mScarlet. Animals were treated as in (A) and 22 animals were imaged over three independent experiments. For each strain, consistent exposure times were used for red and green FP imaging. P values are from a two-tailed Student’s t-test. N.S.=p>0.01; *=p<5×10^14^.

We next tested whether Gamillus affects the function of proteins to which it is fused, using proteins sensitive to tag dimerization. We used CRISPR to knock Gamillus coding sequence into a histone H3B (*his-72*) and lamin *(lmn-1*). We also tagged a germline helicase that localizes to P granules, which are found in ribonucleoprotein condensates. We observed the expected chromatin (*his-72*), nuclear envelope (*lmn-1*), and perinuclear (*glh-1*) localization for each fusion (Fig. 5A-E). Notably, Gamillus∷GLH-1 knock-ins were dimmer than GFP knock-ins (Fig. 5E), consistent with the need to image Gamillus at a wavelength that produces 50% excitation to avoid photoconversion (Fig. 5E; Shinoda *et al*. 2018a). These data suggest that Gamillus does not cause mislocalization and validates the FP for tagging proteins by CRISPR-mediated genome editing. Together, these results validate Gamillus as a green FP option for studying lysosome lumenal proteins.

**Fig 5.**
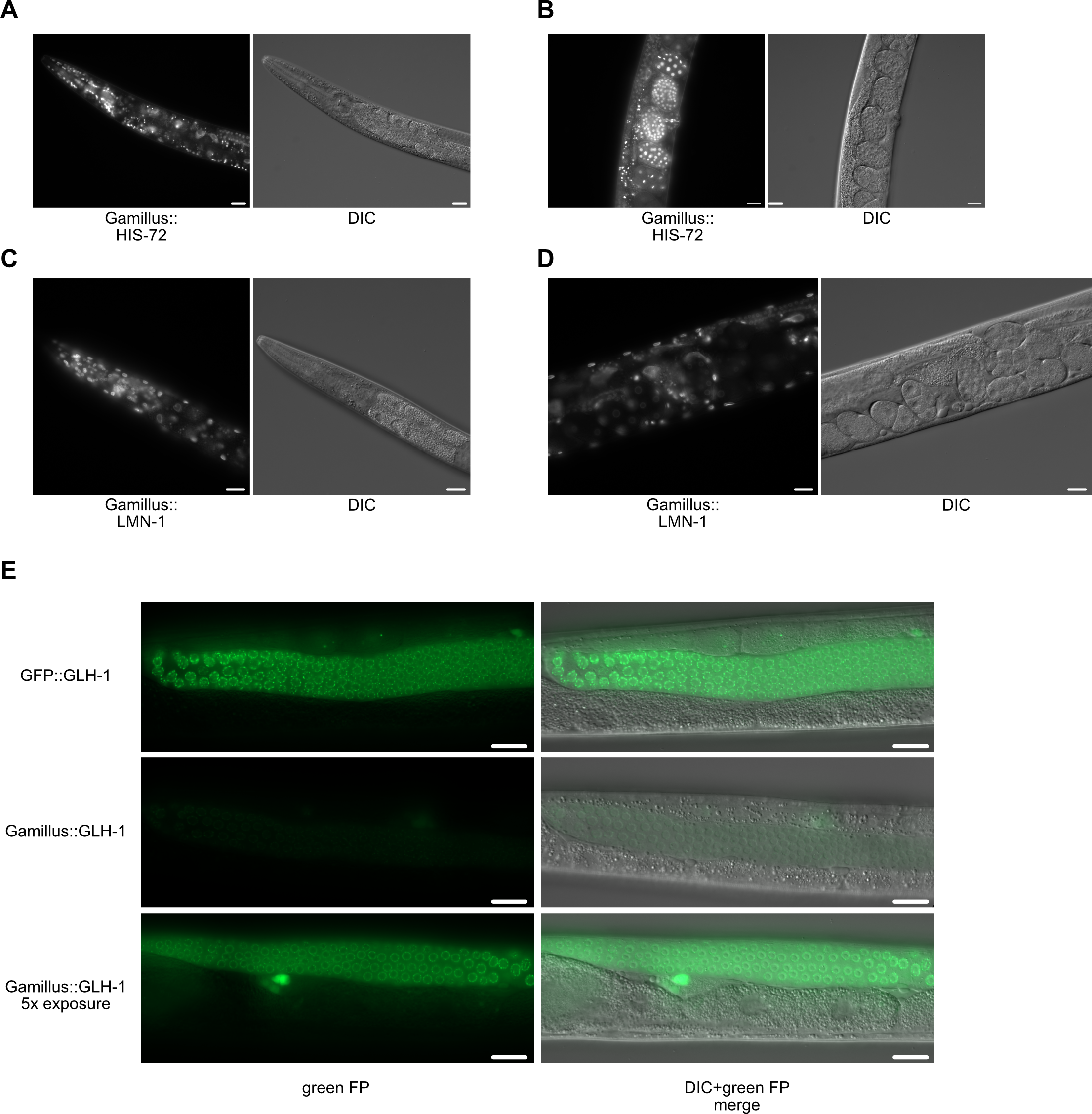
Gamillus does not disrupt localization of *his-72, lmn-1*, and *glh-1* fusion proteins. Gamillus∷HIS-72 and DIC image of adult head (A) and embryos (B). Images are representative of 20 animals from two independent replicates. Scale bars=20 μm. Gamillus∷LMN-1 and DIC image of adult head (C) and embryos (D). Images are representative of 20 animals from two independent replicates. Scale bars=20 μm. € GFP∷GLH-1 and Gamillus∷GLH-1 germline images along with DIC overlays. A Gamillus∷GLH-1 image where a 5x longer exposure was performed is provided. Different animals were imaged for the 1x and 5X Gamillus exposures but all images are representative of 20 animals from two independent experiments. Scale bars=20 μm.

## DISCUSSION

Like our findings with MLT-11 (Fig. 1; Ragle et al. 2022), different localization patterns of green and red FP fusions have been reported for *C. elegans* factors involved in moltin such as MLT-10, NOAH-1, and PTR-4 (Meli *et al*. 2010; Cohen *et al*. 2019, 2021; Johnson *et al*. 2022). Lysosomal localization poses different issues for green and red FP fusions. Green FP degradation and/or quenching could create false negatives for lysosomal localization. Conversely, the stability of red FPs in the lysosome could allow a cleaved red FP tag to accumulate in the absence of the fusion protein, creating a false positive for lysosomal localization of a factor of interest. Additionally, the bright lysosomal signal can produce high background, obscuring dimmer localization of a translational fusion of a protein of interest in other tissues or cellular compartments. Determining the extent of FP cleavage by western blotting is a critical control to interpret any lysosomal localization of fusion proteins. Sample processing made a major difference in MLT-11∷mNG∷3xFLAG degradation during western blotting. While the levels of the full-length protein weren’t obviously affected, the laddering can impair identification of isoforms (Fig. 2). There was a ~50 kDa band that must be produced by C-terminal cleavage of MLT-11 (Fig. 2) for which we have observed a peak in expression early in L4 (Ragle et al., 2022). We are currently pursuing how this isoform is produced and whether it plays any distinct roles in molting. Our data also suggest that FLAG epitopes become unrecognizable by anti-FLAG antibodies after extended time in lysosomal environments (Fig. 3). We observed a similar phenomenon with MLT-11∷mNeonGreen∷3xFLAG where in late-L4 larvae and early adulthood we observed lysosomal localization but no signal by anti-FLAG immunoblotting (Ragle *et al*. 2022). These results are likely due to degradation of the epitope by lysosomal proteases, though we cannot rule out post-translational modification of the FLAG tag in the lysosome that prevents antibody binding. Using antibodies against FPs may be preferable to use in immunoblotting as a way to test whether the fusion protein is full-length. While the rigid proline linker and mCherry N-terminal truncation did not reduce mCherry cleavage from NUC-1, it is possible that they may work on other proteins.

We also validated Gamillus as a green FP for labeling the lysosomal lumen and for fusion to lysosomal proteins. When fused to NUC-1, it displayed similar co-localization and acid-tolerance as mCherry (Fig. 4). We also confirmed that despite its higher pKa than mCherry, mScarlet is acid-tolerant making it suitable for lysosomal experiments (Fig. 4). Gamillus also exhibits photoswitching behavior at its peak excitation wavelength. If non-peak excitation wavelength (440-480 nm) is used the switch to the off-state is minimized at the cost of brightness (Shinoda *et al*. 2018a).

We note that our analyses focused on the hypodermis. Hypodermal lysosomes undergo changes in activity, becoming highly active during the molt when they help recycle cuticular components. Intestinal cells are another major source of lysosomal activity in the animal. Gut granules are specialized lysosome-related organelles that play roles in lipid transfer, metabolism, detoxification, signaling, and zinc storage (McGhee 2007; de Voer et al. 2008). Testing the performance of Gamillus, P5 linkers, and western blot sample processing in this tissue is an important area of future exploration.

## ACKNOWLEDGEMENTS

We thank D. Kellogg for helpful discussions and R.K. Spangler for comments on the manuscript. This work was funded by the National Institutes of Health (NIH) National

Institute of General Medical Sciences (NIGMS) [R01 GM138701] to J.D.W. Some strains were provided by the *Caenorhabditis* Genetics Center, which is funded by the NIH Office of Research Infrastructure Programs [P40 OD010440]. The anti-alpha tubulin 12G10 monoclonal antibody developed by J. Frankel and E.M. Nelson of the University of Iowa was obtained from the Developmental Studies Hybridoma Bank, created by the NICHD of the NIH and maintained at The University of Iowa, Department of Biology, Iowa City, IA 52242.

